# Epistasis and background dependence in the evolution of Omicron variants of the SARS-CoV-2 Spike protein

**DOI:** 10.1101/2025.11.18.689118

**Authors:** Alief Moulana, Thomas Dupic, Michael M. Desai

## Abstract

The rapid and repeated emergence of SARS-CoV-2 variants, particularly within the Omicron lineage, highlights the virus’s remarkable ability to adapt under shifting immune pressures. A central molecular battleground in this evolutionary arms race is the spike receptor-binding domain (RBD), which must simultaneously maintain high affinity for the human ACE2 receptor while evading recognition by neutralizing antibodies. In this study, we construct and analyze multiple combinatorial libraries of SARS-CoV-2 RBD variants spanning major branches of Omicron evolution, including BA.1, BA.2, BA.5, XBB, and JN.1. Using high-throughput yeast display and binding assays, we map the effects of thousands of mutations and their combinations on ACE2 binding and antibody evasion. Our results reveal that while many RBD mutations exhibit additive effects, several mutations interact epistatically in a background-dependent manner. In particular, we identify synergistic interactions between BA.1 and BA.5 mutations that enhance antibody evasion, likely facilitating the rise of recombinant variants and convergent evolution. Conversely, some mutations show lineage-restricted compatibility, suggesting potential constraints on future evolutionary trajectories. Our comprehensive genotype-to-phenotype maps uncover both rugged and smooth regions of the viral fitness landscape and underscore the importance of epistasis in shaping SARS-CoV-2 evolution. These findings improve our ability to anticipate future viral variants and provide a framework for understanding how host-pathogen co-evolution unfolds at the molecular level.

## INTRODUCTION

The SARS-CoV-2 pandemic has disrupted societal norms and placed immense pressure on global healthcare systems, but it has also presented a rare opportunity to observe viral evolution in real time. As the pandemic progressed, the virus continuously evolved in response to shifting population immunity, shaped by vaccination efforts, waves of infection, and public health measures^1^. This ongoing interaction between the virus and host immunity is a prime example of an evolutionary arms race, most evident at the molecular level, where viral antigens and host cell proteins interact.

For SARS-CoV-2, the receptor-binding domain (RBD) of the spike protein plays a central role in this molecular recognition, mediating viral entry through the host ACE2 receptor^2–4^. At the same time, the host’s adaptive immune system mounts a response, primarily through the production of antibodies^5^, which compete for binding to the RBD. This puts the RBD under at least two opposing selective pressures: maintaining high affinity for ACE2 while evading recognition by the immune system. As the virus adapts, each new mutation moves the RBD across a fitness landscape (i.e. a map from amino acid sequences to fitness-relevant phenotypes such as antibody evasion and receptor binding). The effects of mutations on these phenotypes determine the virus’s ability to persist within individual hosts and spread through the broader population^6^.

The initial phase of the pandemic saw the virus rapidly adapting, leading to several common mutations in the receptor-binding domain (RBD). Some mutations (e.g., N501Y) improved binding affinity to ACE2, while others (e.g., K417N and E484K) helped evade antibodies, giving rise to numerous variants of concern (VOCs). By late 2021, as the Delta wave subsided, the Omicron BA.1 variant emerged. Although genetically very distinct, Omicron’s closest relative was the original Wuhan Hu-1 strain and not the then-widespread Delta variant. Compared to Wuhan Hu-1, BA.1 harbors dozens of mutations across its genome, including 31 mutations on the spike protein and 15 within the RBD alone. These figures were considerably higher than those seen in earlier VOCs. This extensive mutational profile allowed the Omicron lineage to escape immunity from prior infections with Delta and other VOCs, as well as existing vaccine-induced immunity^6^.

In previous work, we systematically characterized how combinations of the 15 RBD mutations in Omicron BA.1 affect ACE2 affinity and antibody evasion^7,8^. High-throughput affinity measurements allowed us to estimate the effects of these mutations both individually and in combination^9,10^. We demonstrated that, for immune evasion, one or two escape mutations are generally enough to reduce affinity to distinct classes of neutralizing antibodies, with minimal or no interactions between these mutations. In contrast, mutational effects on ACE2 affinity exhibit pervasive epistasis: while individual escape mutations typically reduce ACE2 binding, these deleterious effects are epistatically compensated by the combination of N501Y and Q498R. This gave the BA.1 variant a substantial fitness advantage over Delta and other pre-Omicron strains, allowing it to spread before being subsequently replaced by the BA.2, BA.5, and recombinant XBB subvariants over the next year of its evolution.

Additional mutations on the Omicron background led to varying immune evasion profiles for each subvariant^11^. In particular, XBB recombinants evaded previous antibodies known to be resistant to other Omicron mutations^12,13^, possibly by combining two lineages that each evade distinct classes of immunity. Little is known, however, about the extent to which these lineage-specific mutations are compatible with each other and what the broader implications for immune evasion are. Further, which selective pressures lead to which diversified branch of the lineage? Could mutations reappear in a totally different branch, and how likely is recombination between branches – do across-branch mutations interact and, if so, how? More broadly, how rugged is the ACE2 landscape and how do the immune landscapes compare?

As this Omicron lineage diversified and novel mutations arose, by the end of 2023 a new subvariant JN.1 completely replaced the other subvariants in a rather similar fashion to its BA.1 ancestor. Like BA.1, JN.1 branched off the BA.2 lineage, accumulating at least 10 additional RBD mutations (comparable to the divergence between BA.1 and Wuhan Hu-1 RBD sequences). However, it is unclear whether JN.1 evolution involves a similar epistatic landscape. Some RBD mutations known to enhance viral fitness have appeared on branches that are distant in time, and previous predictions can sometimes identify the next genotype of interest. But because of potential epistatic effects, our ability to predict phenotypic effects of specific mutations becomes increasingly limited in more distant genetic backgrounds. It is therefore important to assess how widespread these interactions are across the vast RBD genotypic space.

Here, we explore these questions by generating more comprehensive phenotypic measurements across diverse RBD genetic backgrounds. We constructed several spike protein RBD libraries, and use a yeast-display system to measure binding affinities of each variant to the cognate human ACE2 receptor and to a set of monoclonal antibodies. These RBD variant libraries include but are not limited to variants from BA.1 to JN.1. Specifically, we built four variant libraries: (i) all combinations of ACE2 Omicron BA.1 mutations on the Wuhan Hu-1 background (previously published^7,8^); (ii) the entire mutational landscape of BA.1, BA.2, BA.4, and BA.5 on the Omicron background; (iii) combinations of prevalent Omicron mutations on seven divergent genetic backgrounds; (iv) combinations of XBB.1 and JN.1 mutations on four different genetic backgrounds. All these libraries are generated combinatorically, ensuring the appearance of each mutation at least 20 times across the library. These mutations are spread out across the span of RBD, with significant proportion being clustered around the binding motif to ACE2 (Figure 1A). This data provides a basis for improving our ability to characterize fitness-relevant genotype-to-phenotype maps, and thus to understand and predict evolution in this system.

**Figure 1:**
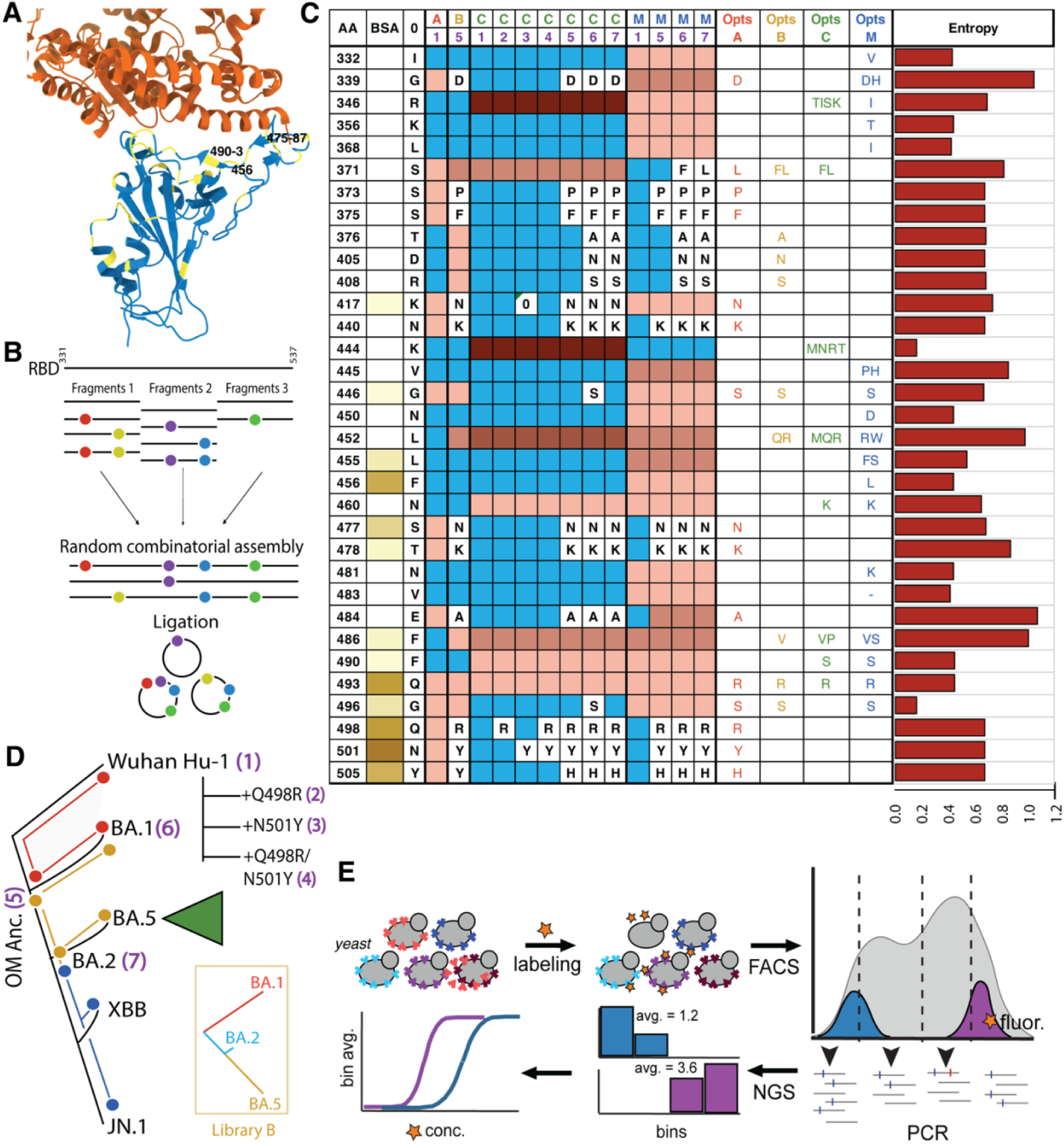
Schematic of variant library design, production, and measurement. (**A**) Co-crystal structure of ACE2 (orange) bound to the RBD (blue), with residues chosen for mutagenesis highlighted (yellow). (**B**) Summary of combinatorial fragment assembly. (**C**) Table summarizing the amino acid positions (y-axis) that are mutated across different libraries (A, B, C, and M columns in various colors) and background combinations (numbered columns in purple). Lettered entries indicate residues that are fixed in a specific library-background combination, while red entries represent polymorphic sites (light to dark red indicating 2 to 4 possible alleles). Blue cells correspond to sites fixed for the Wuhan Hu-1 wildtype residue (column 0). The polymorphic mutation options for each library are listed under Opts A, Opts B, Opts C, and Opts M. Additional site properties include buried surface area (BSA) from the ACE2-bound co-crystal structure and Shannon entropy in natural sequencing data (site variability). (**D**) Evolutionary tree (black line) depicting the relationships among specific variants of concern (nodes). Colored lines indicate the occurrence of mutations included in each variant library along the tree, following the scheme in (**C**) (red: A, orange: B, blue: M), with green triangle representing Library C mutations. Backgrounds used as templates for library construction are numbered in purple corresponding to the columns in (**C**). Inset shows the evolutionary tree between the three main variants included in Library B, with coloring following the scheme in Figure 2. (**E**) Schematic of library transformation into yeast and the subsequent phenotype measurement workflow.

## RESULTS

### Library choice and generation

Our libraries were constructed by combinatorially assembling three to four fragments of the RBD sequence. Each fragment has multiple possible variants (all possible combinations of the mutations within that fragment), and during Golden Gate assembly one version of each fragment is randomly incorporated to assemble a single complete RBD sequence (Figure 1B). In this way, we produce a diverse library containing all (or most) combinations of mutations across the RBD (see Methods and Moulana *et al.,* 2022 for details).

In this study we constructed three yeast-display RBD variant libraries and analyzed them alongside the library we previously constructed in Moulana *et al., 2022.* The fragments for Libraries A, B, and C were created by PCR, using primers with overhangs that contain mutations. The fragments for Library M were synthesized as an oligo pool (Twist Biosciences). Each of these libraries was designed to address a different aspect of RBD evolution during the pandemic (Figure 1C):

i. **Library A** is the original combinatorial library containing all possible combinations of the 15 spike protein RBD mutations that separate Wuhan Hu-1 and Omicron BA.1, previously constructed in Moulana et al., 2022. The library contains 2^15^ = 32,768 variants.
ii. **Library B** spans the entire mutational landscape of BA.1, BA.2, BA.5, and BA.2.12.1, added on the ancestral Omicron background. This involves seven biallelic and two triallelic loci, for a total of 2^7^ x 3^2^ = 1,152 variants. This library focuses on studying how mutations, both within and across lineages, interact to affect ACE2 and antibody binding affinities (Figure 1D).
iii. **Library C** extends our study to include prevalent mutations from the Omicron BA.2 lineage, introduced in all possible combinations on each of seven genetic backgrounds (see Figure 1D), for a total of 7 (backgrounds) x 2^3^ x 3^1^ x 4^1^ x 5^2^ = 16,800 variants. This library investigates how these BA.2 mutations affect ACE2 and antibody affinities across the possible historic backgrounds.
iv. **Library M**, assembled combinatorically from synthesized DNA fragments, contains one of four possible genetic backgrounds (Wuhan Hu-1, Omicron ancestor, BA.1, and BA.2). On each background, combinations of 26 mutations common to the new JN.1 and XBB.1 lineages are introduced. This theoretically results in 4 x 2^16^ x 3^5^ = 63,700,920 variants, of which we randomly produced and assayed ∼300,000.

After constructing each library, we performed various binding assays (Figure 1E; Table 1). These assays broadly measure one of three types of observed phenotypes: absolute binding affinity (K_D_), binding fluorescence at one or two ligand concentrations, and enrichment of fluorescence - labeled binders at a single concentration (see Methods for details). The specific ligand and concentration used for each measurement vary depending on the library (Table 1).

**Table 1:** Library assays. Overview of the high-throughput measurements (ACE2 and mAb) we performed on each library.

| Library | ACE2 Assay | mAb Assay | List of mAbs |
| --- | --- | --- | --- |
| B | TiteSeq | TiteSeq | AZD1061, AZD8895, REGN10987 |
| C | TiteSeq | SortSeq (x2) | AZD1061, AZD8895, REGN10987, LY-CoV1404, S309 |
| M | SortSeq | Enrichment | AZD1061, AZD8895, REGN10987, LY-CoV1404, S309, 002-S21F2, Iv0221, Iv0262 |

Across the three libraries, we measured affinities to the class 1 antibody AZD8895 and the class 3 antibodies AZD1061 and REGN10987. These two classes target distinct RBD epitopes^14,15^ and differ in their efficacy against the two main Omicron clades, BA.1 (still susceptible to class 1) and BA.2 (still susceptible to class 3)^11,16^. For Library C, we additionally measured binding to LY-CoV1404, a class 3 antibody that retains binding against most BA.2 subvariants, including BA.5, but is evaded by some common BA.5 mutations^17,18^. For Library M, we further included three class 3 antibodies (S309, 002-S21F2, and Iv0221), which still bind some of the later BA.2 subvariants^19,20^.

These datasets demonstrate strong reproducibility across assay types. The K_D_ measurements for Library B are strongly correlated between replicates (Supplementary Figure 1A; R² > 0.91). Similarly, fluorescence and enrichment measurements in subsequent libraries also exhibit high correlations (Supplementary Figure 1B-C; R² > 0.88 and R² > 0.76, respectively).

### Beneficial combinations of lineage-specific mutations

Omicron BA.1, BA.2, and BA.5 each carry lineage-defining mutations (all included in Library B), with each lineage affecting distinct sets of measured phenotypes (Figure 2A). Specifically, BA.1 mutations confer REGN10987 evasion, whereas BA.5 mutations affect ACE2 and AZD8895 binding affinities. In contrast, AZD1061 affinity appears to be similarly affected by both lineages. Interestingly, BA.2 mutations show little contribution to either enhanced ACE2 affinity or antibody evasion in our assays. Therefore, BA.1 and BA.5 mutations appear to drive Omicron lineage evolution, while BA.2 mutations may either be largely neutral or potentiate later adaptation.

**Figure 2:**
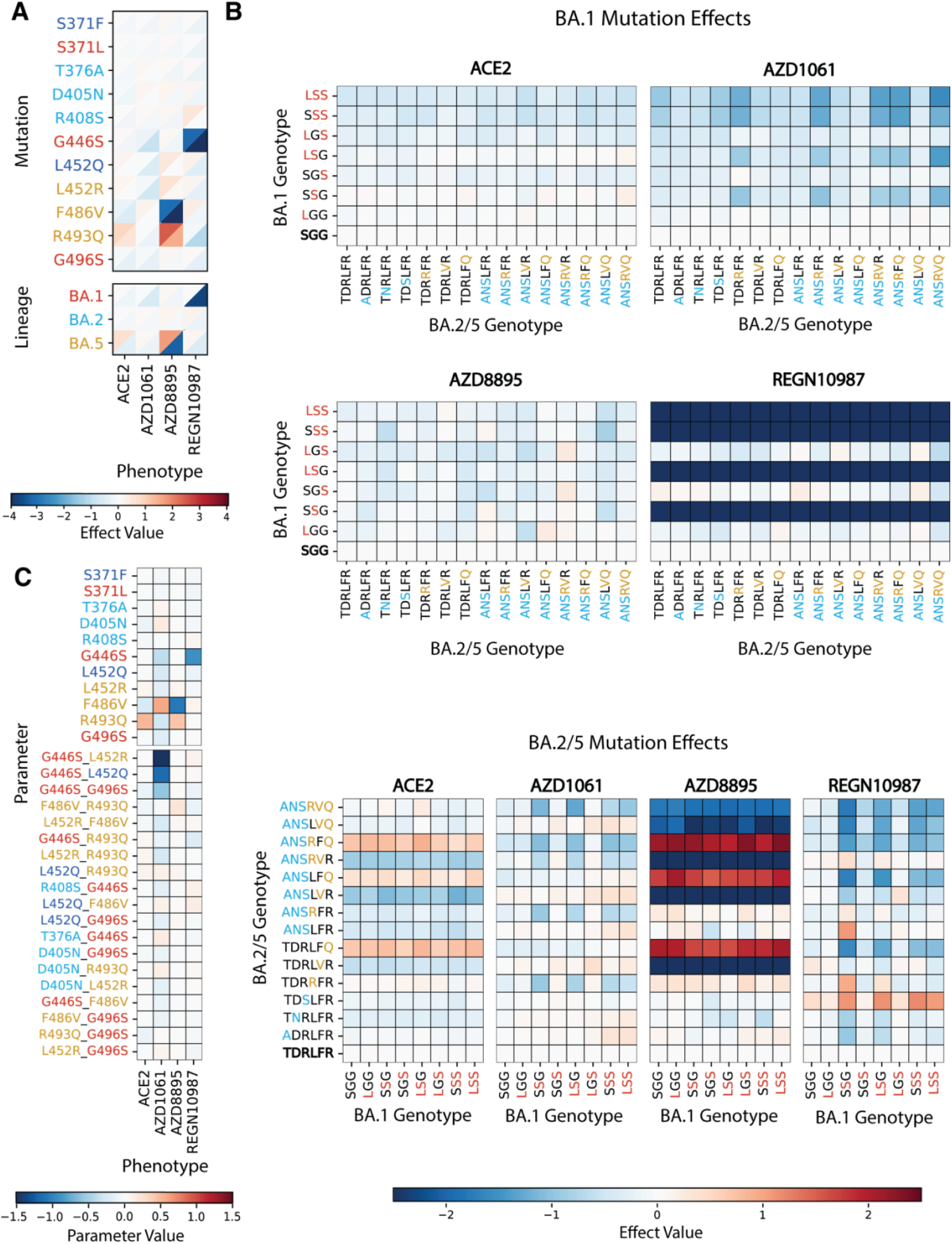
Combinations of lineage-specific mutations. (**A**) Heatmaps showing the single-mutation effect values (color scale: low → high, blue → red), across all mutational backgrounds for each binding affinity (-log *K*_D,app_) phenotype (x-axis). The upper left triangle indicates the 75th percentile of mutation effects, while the lower right triangle represents the 25th percentile. Single-mutation effects are grouped by the viral lineage where the mutation naturally occurs (BA.1, BA.2, BA.5), with cumulative effect for each background displayed below the single-site effects. Mutation names are colored according to this grouping: red for BA.1 mutations (S371L, G446S, G496S), light blue for mutations shared between BA.2 and BA.5 (T376A, D405N, R408S), orange BA.5-specific mutations (L452R, F486V, R493Q), and dark blue for other mutations (S371F and L452Q). (**B**) Heatmaps showing combined effects of BA.1 mutations across genotypes carrying different BA.2/5 mutations (top) and of BA.2/5 mutations across genotypes carrying different BA.1 mutations (bottom), excluding mutations S371F and L452Q. Sites are colored according to the scheme in (**A**), with the unmutated allele being black. Effects are computed for each column (x-axis; genotype mutations appear on), where the phenotype of the variant in each row is normalized by subtracting the phenotype of the unmutated variant (bottom row). (**C**) Heatmaps of inferred model parameters for single and pairwise mutations, evaluated across different phenotypes for models truncated at order 2. Pairwise interactions with a sum of absolute values exceeding 0.25 are shown, where each mutation pair is separated by a dash. Parameter names are colored based on scheme in (**A**).

In general, Library B variants bind ACE2 with higher affinity and monoclonal antibodies with lower affinity than the Wuhan Hu-1 variant (Supplementary Figure 2). This suggests that most combinations of BA.1 and BA.5 mutations do not substantially reduce fitness. Additionally, about one-third of the library binds ACE2 with even higher affinity than BA.1, BA.2, and BA.5 variants, and a large proportion of the library also shows reduced affinity compared to these variants of concerns. Thus, some alternative mutation combinations may actually improve fitness relative to established variants. Indeed, the addition of further mutations on the Omicron ancestral genotype does not decrease the average ACE2 affinity while simultaneously reducing binding to the monoclonal antibodies (Supplementary Figure 3). This pattern may result from the accumulation of strongly beneficial mutations or from potential synergistic epistasis between mutations from different lineages. The latter is particularly relevant given the possibility that recombinants between distinct lineages could emerge with higher-than-additive fitness.

Such synergetic epistasis between BA.1 and BA.2/5 mutations are not observed for both ACE2 and AZD8895 affinity (Figure 2B). Although adding combinations of BA.5 mutations, especially F486V and R493Q, do affect ACE2 affinity, these effects are independent of additional BA.1 mutations. Additionally, our epistasis model only detects significant first-order effects and not pairwise ones (Figure 2C). Similarly, for AZD8895, the two BA.5 mutations do affect affinity, but their effects are generally consistent, with minimum interactions. Overall, BA.5 mutations alone affect these two phenotypes, without any observed interactions with BA.1 mutations or among the BA.5 mutations themselves.

In contrast, AZD1061 and REGN10987 affinity landscapes suggest potential beneficial interactions between mutations fixed in distinct lineages. Notably, BA.5 mutations confer greater immune escape in the presence of the BA.1 mutation G446S, while G446S itself exhibits stronger effects on backgrounds containing the BA.5 mutation L452R (Figure 2B). Indeed, our epistasis model identifies only two significant interactions across all phenotypes, G446S and L452R/Q, both affecting AZD1061 affinity (Figure 2C). In comparison, on REGN10987, interaction between G446S and BA.5 mutations appears relatively minor compared to the large single-mutation effect of G446S (Figure 2B,C). Consistent with Library A results, Omicron BA.1 variant shows no detectable binding to REGN10987. Nevertheless, BA.5 mutation R493Q does reduce REGN10987 affinity, particularly on genotypes containing G446S. Altogether, our Library B dataset shows that BA.1- and BA.5-specific mutations affect distinct subsets of phenotypes, yet for AZD1061 and REGN1098 evasion, BA.5 mutations can interact synergistically with the BA.1 mutation G446S.

### BA.5 mutations have lineage-specific effects

After the BA.5 lineage rose to dominance, several convergent mutations began to accumulate across its subclades, many of which further improve variant fitness within the lineage. Here, we constructed combinatorial Library C by introducing a set of convergent mutations not only onto the BA.5 lineage itself but also across several variant backgrounds. Many of the resulting combinatorial variants exhibit reduced affinity to monoclonal antibodies, particularly compared to previously established variants (Supplementary Figure 4). Nevertheless, these variants largely retain ACE2 binding, with some even displaying higher affinity than the established variants.

Individually, the convergent mutations across the BA.5 clade have little effect on ACE2 affinity (Figure 3A). Instead, in Library C, much of the observed variance in ACE2 binding can be explained by the underlying genetic backgrounds. Consistent with Library A results, the compensatory mutations Q498R and N501Y (Background 4) improve ACE2 affinity. As seen in Library B, the BA.1- and BA.2-specific mutations (Backgrounds 6–7) slightly reduce ACE2 affinity compared to the ancestral Omicron background (Background 5), yet their ACE2 affinities remain higher than that of the Wuhan Hu-1 variant (Background 1). In contrast, BA.5 mutation themselves have relatively weaker effects on ACE2 affinity. Although substitutions at RBD sites 486 and 493 generally reduce binding, these effects are mitigated in BA.5 through two mechanisms: (i) the F486V/S substitutions are less deleterious on Omicron-derived backgrounds (Figure 3B; Backgrounds 5–7), and (ii) the Q493R substitution is largely replaced by its reversion, R493Q. Therefore, among the common BA.5 mutations, only the fixed substitutions R493Q and F486V contribute to maintaining ACE2 affinity.

**Figure 3:**
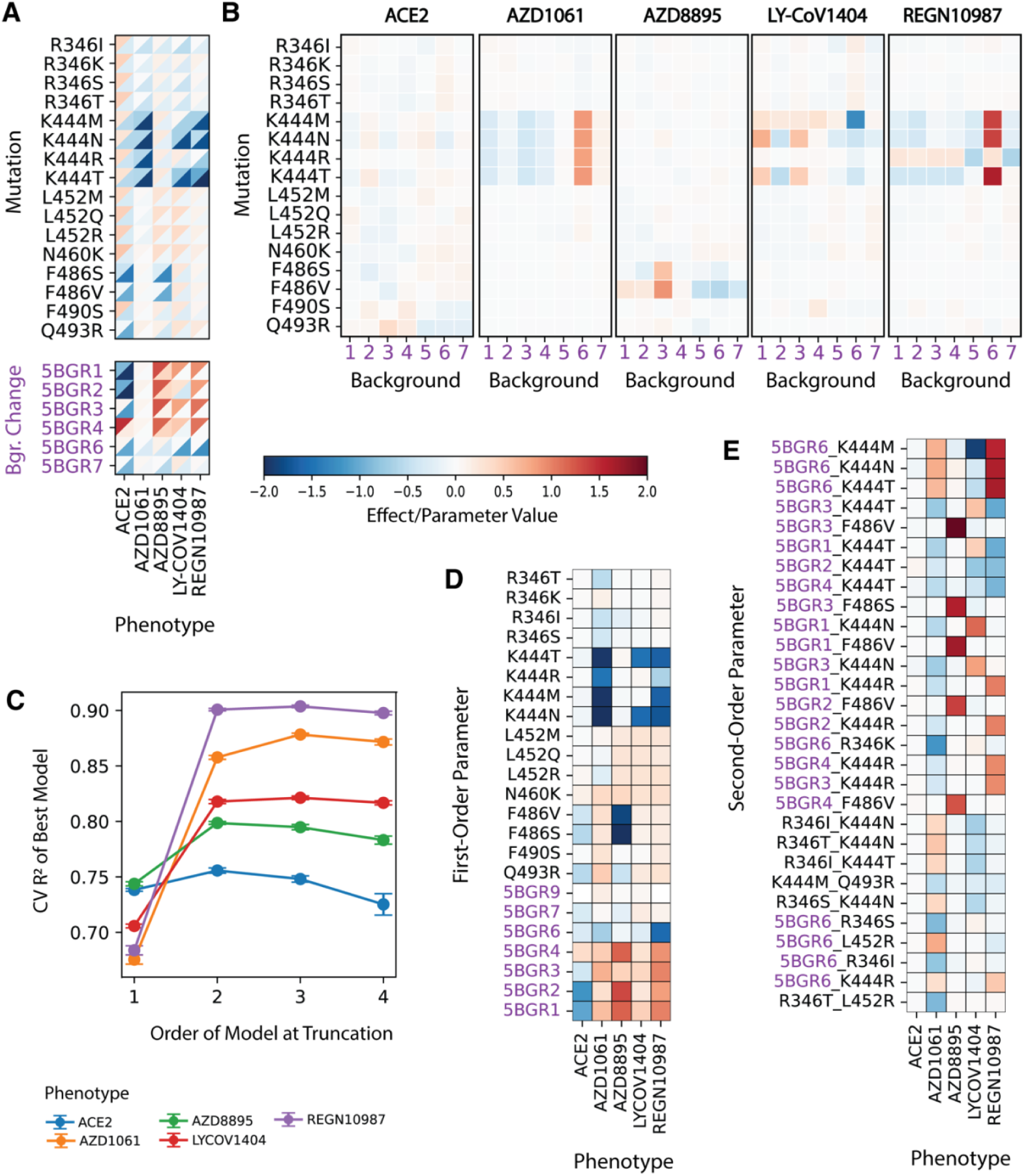
BA.5 mutations across different genetic backgrounds. (**A**) Heatmaps showing the single-mutation effect values (color scale: low → high, blue → red) across all mutational backgrounds for each phenotype (x-axis). The upper left triangle indicates the 75th percentile, while the lower right triangle represents the 25th percentile of mutational effects. Combined effects of mutations in each background (in purple, relative to background 5, ancestral Omicron) are shown below the single-site effects. (**B**) Heatmaps displaying the mean effect of each mutation (y-axis) across different backgrounds (x-axis) for each assayed phenotype. To emphasize background-specific effects, the mean across backgrounds is subtracted from each row, effectively mean-centering the data. (**C**) Cross-validated performance (R^2^) for each model across 10-fold cross-validation. Points indicate the mean (R^2^) across folds, with error bars representing standard deviation. (**D,E**) Heatmaps of inferred model parameters for single (**D**) and pairwise (**E**) mutations, evaluated across different phenotypes for models truncated at order 3. Pairwise interactions with a sum of absolute values exceeding 1 are shown, where each mutation separated by a dash. Background changes are included as one parameter, colored in purple.

In contrast, common BA.5 mutations strongly promote antibody evasion, with substitutions at RBD site K444 being particularly effective against all Class 3 antibodies tested: AZD1061, LY-CoV1404, and REGN10987 (Figure 3A). However, the effects of K444 and other BA.5 mutations differ across genetic backgrounds (Figure 3B), as also reflected in the epistasis model predictions (Figure 3C–E). For ACE2 affinity, including higher-order interaction terms only marginally improves or even reduces model performance (Figure 3C), whereas pairwise terms markedly improve predictions for antibody-binding phenotypes. These apparent interactions may reflect true epistasis between mutation and background or limits of measurement sensitivity. For example, only two K444 alleles (T and N) confer escape from LY-CoV1404, while K444M mediates escape only on the BA.1 background (Background 6), indicating indeed an interaction between this site and BA.1 mutations (Figure 3B,E). In contrast, for AZD1061 and REGN10987, BA.1 mutations themselves already reduce binding relative to the Omicron ancestor (Figure 3A,D), which likely dampen the apparent impact of K444 mutations on this background (Figure 3B,E). This dampening is particularly evident for REGN10987, which is effectively escaped by the BA.1 variant, as previously observed^8^. Altogether, although BA.1 mutations alone are already effective at evading Class 3 antibodies, particularly REGN10987 and AZD1061, the acquisition of K444 mutation is either sufficient or necessary (as in the case of LY-CoV1404) to achieve broader Class 3 antibody escape.

Although not to the extent of the strong K444 mutations, other BA.5 lineage substitutions still affect antibody escape, sometimes with background dependence. As observed in Library B, F486V/S variants effectively escape the class 1 antibody AZD8895, which is not escaped by K444. These effects are less pronounced in pre-Omicron backgrounds (Backgrounds 1–4; Figure 3B), suggesting that acquisition of F486V/S in BA.5 specifically contributes to AZD8895 or similar antibody escape. R346 mutations, in contrast, contribute to AZD1061 escape with minimal background dependence. R346T interacts with L452R to produce stronger evasion effects (Figure 3E) and also interacts with K444N/T for enhanced LY-CoV1404 escape. In fact, such interactions between two BA.5 mutations are much rarer than those involving a background change (Figure 3E), indicating that the landscape is largely shaped by background dependence rather than interactions among BA.5-specific mutations themselves.

While phenotypic landscapes spanned by Library C are largely additive (Figure 3C-E), the specific compatibility of BA.5 mutations within their native lineage background suggests that this type of interactions may have influenced their evolutionary trajectory. However, this background dependency, along with the diverging effects of individual mutations across different phenotypes, may have constrained BA.5 evolution. Although BA.5 experienced a rapid expansion, its dominance was short-lived, as it was soon displaced by BA.2 recombinants, such as XBB, and later by JN.1. These patterns highlight the dynamic nature of SARS-CoV-2 evolution, where the fitness landscape continuously shifts due to both mutation interactions and immune-driven selection pressures.

### JN.1 and XBB mutations slightly prefer BA.2 lineage

The dominance of the BA.5 lineage waned following the reemergence of BA.2-derived recombinant lineages, notably XBB, a recombinant between BA.1 and BA.2, and subsequently JN.1. These lineages harbor distinct yet overlapping sets of prevalent mutations relative to their ancestral BA.2 background (Figure 1B-C; Figure 4). To investigate the phenotypic impacts of these mutations, we generated variant libraries consisting of random combinations of JN.1 and XBB mutations introduced onto four representative genetic backgrounds (Wuhan Hu-1, ancestral Omicron, BA.1, and BA.2; Table 1). Consequently, our data provide statistical summaries of phenotype effects for each mutation within these distinct genetic backgrounds rather than exhaustive measurements on each possible combinatorial background, as was done for Libraries A, B, and C (see Methods).

**Figure 4:**
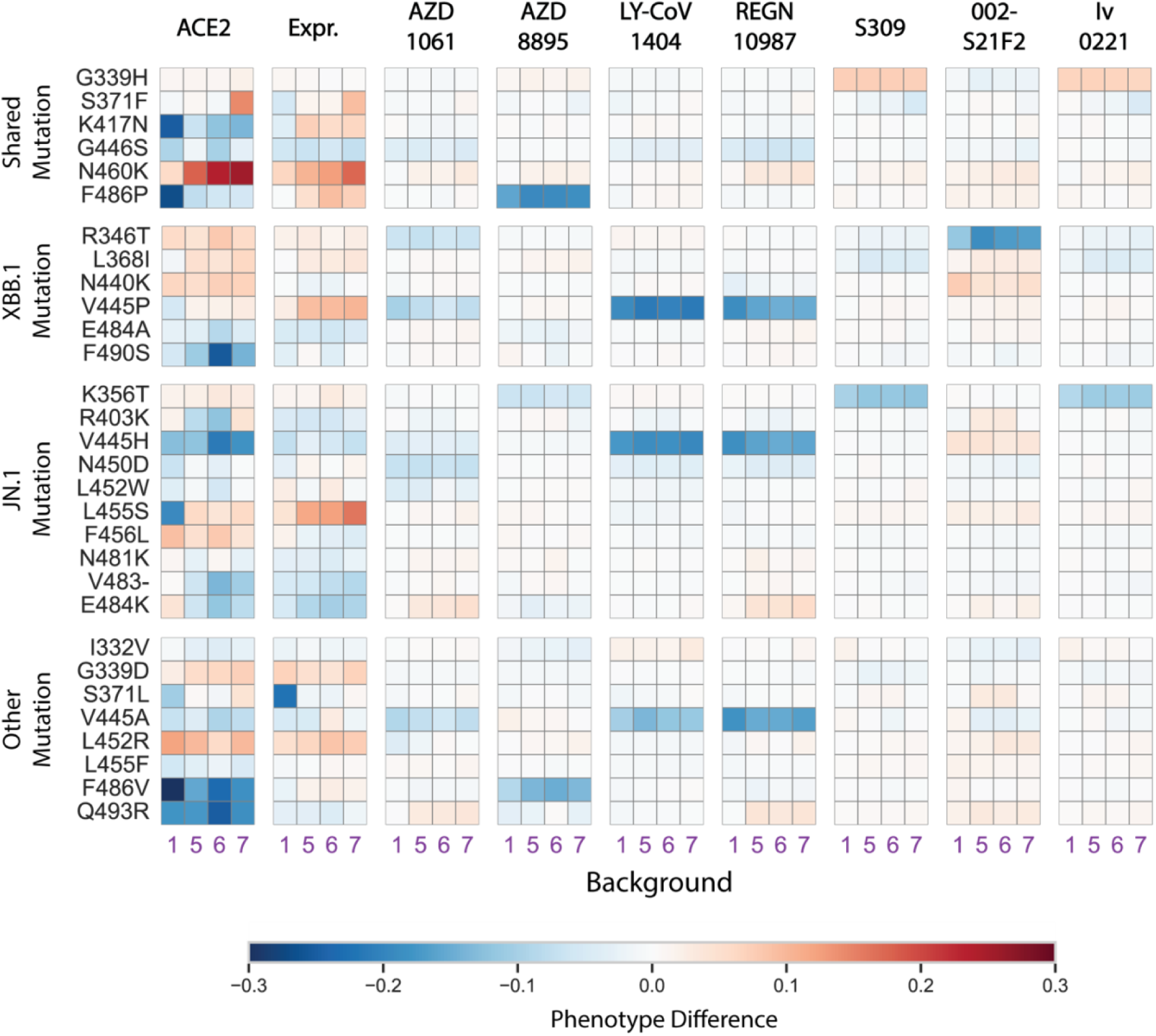
XBB and JN.1 mutations across different genetic backgrounds. Heatmaps showing the mean effect of each mutation (y-axis) across different genetic backgrounds (x-axis) for each measured phenotype. For each background, the effect of a mutation is calculated as the difference between the mean phenotype of genotypes carrying the mutation and the mean phenotype of genotypes lacking it. Mutations are grouped by their occurrence in both lineages, only the XBB lineage, only the JN.1 lineage, or other BA.2 lineages.

We find that the effects of XBB.1 and JN.1 mutations on ACE2 affinity and protein expression depend strongly on genetic background (Figure 4). Five mutations shared between the two lineages (N460K, S371F, F486P, G446S, and K417N) substantially alter ACE2 binding, but their effects are highly background dependent. N460K and S371F improve ACE2 affinity on Omicron-derived (5–7) backgrounds but have little impact on the ancestral Wuhan Hu-1 (1) background. Conversely, F486P, G446S, and K417N are more deleterious on BA.1 (6) and Wuhan Hu-1 (1) backgrounds. Together, these patterns indicate a modest preference for BA.2 (7) and ancestral Omicron (5) backgrounds. Expression levels show a similar trend, with stronger beneficial effects on BA.2 and other Omicron-derived contexts. The same background dependency extends to lineage-specific substitutions: for example, JN.1-specific mutations R403K and L455S enhance ACE2 binding on the BA.2 background despite being deleterious elsewhere. Overall, background-dependent effects dominate the phenotypic variance in ACE2 affinity and are generally stronger than any detectable pairwise mutation interactions (Supplementary Figure 5).

In contrast to their effects on ACE2 affinity, the effects of these mutations on antibody evasion are more consistent across genetic backgrounds. Most shared XBB.1 and JN.1 mutations do not confer substantial escape, except for F486P, which reduces binding to AZD8895. Evasion from LY-CoV1404 and REGN10987 arises from distinct substitutions at the same residues in each lineage, suggesting convergent solutions to immune pressure. Notably, the last three antibodies tested show lineage-restricted escape. R346T, unique to XBB.1, strongly reduces binding by 002-S21F2, an antibody known to neutralize both BA.1 and BA.2^20^. Meanwhile, K356T, specific to JN.1, mediates escape from S309 and Iv0221—broadly neutralizing antibodies that remain effective against all prior Omicron variants. The partial evasion of these antibodies by K356T underscores the capacity of JN.1 and its descendants to further erode broad antibody protection. While most evasion phenotypes appear additive, especially with backgrounds, several substitutions exhibit measurable epistasis, sometimes exceeding background effects (Supplementary Figure 5). Detecting higher-order interactions, however, will require substantially larger datasets and more precise assays.

Together, these findings suggest that the emergence of the XBB.1 and JN.1 lineages reflects a balance between maintaining ACE2 affinity and broadening immune evasion. Several key escape mutations (e.g., V445P, V445H, F486P) reduce ACE2 affinity, but these deleterious effects are mitigated by compensatory substitutions such as N460K and S371F, both of which are more beneficial on the Omicron BA.2 background. Similarly, across Libraries C and M, mutation effects tend to be more beneficial or less deleterious on Omicron-derived backgrounds (Figures 3–4). However, among themselves, post-Omicron mutations are largely additive, with little evidence of strong epistasis, particularly for ACE2 affinity. This contrasts with Library A results, where the ACE2 landscape appears more rugged than those of antibody escapes. Consequently, the phenotypic landscapes following initial Omicron diversification seem relatively smooth, with most interactions confined to those between new and ancestral Omicron mutations rather than among the new mutations themselves. This may help explain why adaptive substitutions continued to accumulate within Omicron-derived lineages with little constraint, as the lineage itself appears to potentiate the acquisition of new mutations compared to its Wuhan Hu-1 predecessor.

## DISCUSSION

Throughout its evolutionary history, SARS-CoV-2 has continuously acquired mutations, particularly concentrated within the receptor-binding domain (RBD) of its spike protein. The phenotypic effects of these mutations can vary substantially depending on the genetic background at other loci, often resulting in complex interactions^7,21^. Such epistatic interactions can enable certain mutations to compensate for otherwise deleterious effects on distinct phenotypes^7^. In this study, we systematically explored this complexity by characterizing multiple genotype spaces across different phenotypic traits. Specifically, we constructed multiple combinatorial libraries reflecting diverse evolutionary branches of the Omicron lineage and measured several binding phenotypes using high-throughput experimental assays.

In the evolutionary history of the Omicron variant, the initial BA.1 variant was rapidly replaced by subsequent BA.2 and BA.5 variants, likely reflecting the higher relative fitness of the BA.5 variant given immune pressures at the time^6^. However, the differential antibody-escape profiles observed between BA.1 and BA.5 variants^22,23^ potentially suggest distinct selective pressures acting upon each subvariant. Notably, BA.5 demonstrates broader antibody evasion compared to BA.1 and BA.2, including resistance to immunity induced by previous infections with these earlier Omicron subvariants^24,25^. Despite these differences, we found beneficial mutations unique to each lineage (e.g., BA.1 and BA.5) to be not necessarily incompatible with each other. Therefore, mutations from both lineages may combine and produce variants of even greater concern. More strikingly, we observed that mutations from distinct lineages may even interact synergistically (Figure 2). For instance, the BA.1 mutation G446S shows positive epistasis with BA.5 mutations. Such interactions may have facilitated the emergence and subsequent spread of recombinant variants, particularly those carrying G446S, following the dominance of BA.5^26,27^.

Within the BA.5 lineage itself, several variants emerged carrying mutations that were not initially fixed but later converged across independent evolutionary branches^28,29^. This diversification within BA.5 may have been driven by heterogeneity in population-level immunity^30,31^. In our analysis of Library C, which encompassed combinations of some of these mutations, we found that mutation effects were typically independent of genetic background. Nevertheless, in certain cases, specific mutations exerted stronger beneficial effects on the BA.2 (BA.5’s progenitor) genetic background, such as mutations at position 444. The specific combination of K444T, N460K, and R346T defines the BQ.1.1 variant, which became a prominent variant of concern primarily due to its potent evasion of class 3 antibodies^32^. Following this pattern, BA.5 was eventually displaced, first by the XBB variant and subsequently by JN.1, potentially reflecting the repetition of the initial Omicron evolutionary dynamics.

The subsequent evolution of post-Omicron lineages was characterized by the accumulation of numerous mutations across different BA.2-derived clades. As a result, unlike in pre-Omicron lineages, many distinct combinations of mutations could give rise to potentially fit variants. One such example is the XBB lineage, a recombinant between two BA.2 descendants, BJ.1 and BM.1.1, which rapidly rose to dominance. The recombination breakpoint lies within the RBD, where each parental segment contributes escape from distinct antibody profiles^33,34^. Following the dominance of the XBB clade, the JN.1 lineage emerged, evolving from the older BA.2-derived subvariant BA.2.86^35^. One notable mutation unique to this clade is L455S, which was found to enhance ACE2 affinity^36^. Interestingly, we found that this effect becomes deleterious on the Wuhan Hu-1 background (Figure 4). More generally, for ACE2 landscape, interactions between the examined mutations and backgrounds are more prevalent than interactions among the mutations themselves. Conversely, weak inter-mutation interactions do appear in the antibody-evasion landscapes, but their effect sizes are relatively small compared to those of single mutations (Supplementary Figure 5). Thus, the explosion of BA.2 derivatives is not likely to have resulted from significant interaction between these novel mutations but might owe to the interaction with the older Omicron ancestral mutations. As a result, different branches could co-occur, and new variants stemming from older clades might emerge, which would have been unlikely on the pre-Omicron clades.

Leveraging our high-throughput phenotype measurements, we have generated detailed genotype-to-phenotype maps for several regions of the SARS-CoV-2 RBD mutational landscape. Specifically, our results highlight scenarios in which viral evolution follows primarily additive landscapes and other cases where it traverses more epistatic spaces. Nonetheless, our current analysis is limited to mutations within the RBD region and thus cannot account for potential epistatic interactions with other protein domains. Additionally, our assays assessed up to three hundred thousand variants, which, although extensive, still represent only a fraction of the vast combinatorial mutation space. More comprehensive datasets that cover additional mutation combinations, broader phenotypes, and more precise measurements are needed to enhance the predictive accuracy and generalizability of evolutionary models. Moving forward, these methods can be further utilized not only to refine predictive genotype-to-phenotype models but also to provide critical insights into the dynamics underlying viral evolution itself.

## METHODS

### Yeast display plasmid, strains, and library production

To generate clonal yeast strains, we used previously generated pETcon-based plasmids expressing Wuhan Hu-1 (pOM102; pAB101), Omicron ancestor (pOM104; pAB105), and Omicron BA.1 (pOM101; pAB106) RBDs. In addition to the RBD coding sequence, which is fused to Aga2 and tagged with myc, this plasmid also includes ampicillin resistance cassette for bacterial selection and a yeast mating factor for protein localization. Using site-directed mutagenesis (NEB #E0554S), we generated additional plasmids expressing RBD variants Wuhan Hu-1 + Q498R (pAB102), Wuhan Hu-1 + N501Y (pAB103), and Wuhan Hu-1 + Q498R + N501Y (pAB104). We also produced an Omicron BA.2 construct via Gibson assembly, replacing the RBD sequence with a gBlock (IDT DNA). All plasmids were amplified in NEB 10-Beta cells and extracted from saturated cultures (Geneaid #PD300). We then transformed Sanger-verified plasmids into the AWY101 yeast strain^37^ as described in Gietz and Schiestl^38^.

To construct RBD variant libraries, we used a Golden Gate combinatorial assembly strategy. Full-length RBD sequences were assembled from sets of dsDNA fragments of roughly equal size, with each set containing fragment versions that differed by the mutations included. For Library B and Library X, fragments were amplified from corresponding plasmids (pAB104 for Library B; pARB101–107 for Library X) using primers introducing the desired mutations and flanked by BsaI recognition sites (Supplementary File 1). For Library M, we produced four fragments: the first two incorporated all combinations of mutations shown in Figure 1A, while the last two included randomly selected combinatorial mutations (Python randomization). We ordered oligo pools (TWIST Bioscience) containing the additional BsaI sites and distinct flanking sequences for each fragment and genetic background. Separate Golden Gate assembly reactions were performed for each library, generating between 2.4 and 10 million E. coli colonies after transformation and recovery in 100 mL of molten LB (1% tryptone, 0.5% yeast extract, 1% NaCl) containing 0.3% SeaPrep agarose (VWR, Radnor, PA #12001– 922) with ampicillin, spread into a thin layer (about 1 cm deep). We then extracted plasmid DNA from the pooled colonies, transformed the libraries into the AWY100 yeast strain, and recovered them in 100 mL of molten SDCAA [6.71 g/L YNB without amino acids (Sigma-Aldrich #Y0626), 2% dextrose (VWR #90000–904), 5 g/L Bacto casamino acids (VWR #223050)], containing 0.5% SeaPrep agarose. From these, we inoculated in liquid SDCAA and froze libraries containing approximately one million yeast colonies each.

### High-throughput binding affinity assays (TiteSeq, SortSeq, and Enrichment)

For some measurements (see Table 1), we performed Tite-seq assays as previously described^9,10,39,40^, with two replicates for each antibody or ACE2. Briefly, we thawed a given yeast RBD library by inoculating the corresponding glycerol stocks in SDCAA (6.7 g/L YNB without amino acid [VWR #90004-150], 5 g/L ammonium sulfate [Sigma-Aldrich #A4418], 2% dextrose [VWR #90000–904], 5 g/L Bacto casamino acids [VWR #223050], 1.065 g/L MES buffer [Cayman Chemical, Ann Arbor, MI, #70310], 100 g/L ampicillin [VWR #V0339]) at 30°C for 20 hr. The cultures were then induced in SGDCAA (6.7 g/L YNB without amino acid [VWR #90004-150], 5 g/L ammonium sulfate [Sigma-Aldrich #A4418], 2% galactose [Sigma-Aldrich #G0625], 0.1% dextrose [VWR #90000–904], 5 g/L Bacto casamino acids [VWR #223050], 1.065 g/L MES buffer [Cayman Chemical, Ann Arbor, MI, #70310], 100 g/L ampicillin [VWR # V0339]), and rotated at room temperature for 18 hr.

Following overnight induction, we pelleted, washed (with 0.01% PBSA [VWR #45001–130; GoldBio, St. Louis, MO, #A-420–50]), and incubated the cultures with monoclonal antibody or ACE2 at corresponding concentrations. The yeast-antibody mixtures were incubated at room temperature for 20 hr. The cultures were then pelleted washed twice with PBSA and subsequently labeled with PE-conjugated goat anti-human IgG (1:100, Jackson ImmunoResearch Labs #109-115-098; for antibodies) or PE-conjugated streptavidin (1:100, ThermoFisher #12-4317; for RBD) and FITC-conjugated chicken anti-cMmyc (1:100, Immunology Consultants Laboratory Inc., Portland, OR, #CMYC-45F). The mixtures were rotated at 4°C for 40 min and then washed twice in 0.01% PBSA.

After incubation and washing, we sorted cells on a BD FACS Aria Illu. Cells were gated by FSC vs SSC and then by expression (FITC) and/or binding fluorescence (PE). The machine was equipped with 405 nm, 440 nm, 488 nm, 561 nm, and 635 nm lasers, and an 85 micron fixed nozzle. Note that, prior to further quantitative assays, we enriched Library M for ACE2 binders (PE+ population) by sorting, recovering, and freezing 8 million ACE2 binding cells. This sorted population is then used for subsequent assays for Library M.

In the **TiteSeq** and **SortSeq** assays, FITC gates were first drawn, and FITC+ populations are sorted into four PE (binding) gates, where the first gate is PE- (non-binding) subpopulation, and PE2, PE3, and PE4 are three PE+ gates with equal population size. Both TiteSeq and SortSeq assays are conducted very similarly, where TiteSeq assay uses a titration framework with many concentrations (10^-12^ to 10^-6^ M at 0.5 log interval) and SortSeq assay on a few concentrations (two concentrations for Library C antibody assay, 10^-6^ and 10^-8^ M, or one concentration for Library M ACE2 assay, 10^-7^ M). In particular, SortSeq allows us to maximize the number of antibodies used (Library C) and the number of cells sorted (Library M). In contrast, **enrichment** assays for antibodies on Library M sort cells into one, instead of four or more, populations. Here, a single PE/FITC gate is drawn, where either non-binding or binding cells are collected, whichever is the smaller population.

In total, for each sample (i.e., independent concentration) across gates, we sorted ∼200,000 yeast cells from Library B, ∼1 million yeast cells from Library C, and ∼4 to 9 million yeast cells from Library M. Sorted cells were then pelleted, resuspended in SDCAA, and rotated at 30℃ until late-log phase (OD600 = 0.9-1.4). The cultures were then pelleted and stored at-20℃ for at least twelve hours prior to extraction using Zymo Yeast Plasmid Miniprep II (Zymo Research # D2004), following the manufacturer’s protocol. The sequencing amplicon libraries were then prepared by a two-step PCR as previously described^10,40,41^.

In brief, we added to the amplicon unique molecular identifies (UMI), inline indices, and partial Illumina adapters through a 7-cycle PCR which amplifies the RBD sequence in the plasmid (Supplementary File 2). We then used the cleaned product (using 0.9X Aline beads) from the first PCR in the second PCR to append Illumina i5 and i7 indices accordingly. The products were then cleaned using 0.85X Aline beads, verified using 1% agarose gel, quantified on Spectramax i3, pooled, and verified on Tapestation 5000HS and 1000HS. Final library was quantitated by Qubit fluorometer and sequenced on Illumina Novaseq SP, supplemented with 10% PhiX.

### Sequence data processing

Following Moulana and Dupic et al.^7,8^, we processed raw demultiplexed sequencing reads to identify and extract the indexes and mutational sites. Briefly, for each antibody, we parse through all fastq files and group the reads according to inline indices, UMIs, and sequence reads. We converted accepted sequences into corresponding nucleotide sequences using regular expression with some amount of mismatch tolerance throughout the RBD sequence. Reads containing errors at known mutation sites were removed. Then for each recovered sequence, counts in each sample were estimated by number of UMIs from all samples into a single table.

### Enrichment inference

For enrichment assays, we calculated an enrichment score. For each variant, we estimated the proportion of cells in the sorted population relative to the total population, 𝑝_𝑠,𝑓_, as follows:

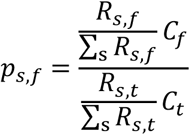

where 𝑅_𝑠,𝑓_ is the number of sequencing reads corresponding to genotype *s* in the fluorescently sorted population, and 𝑅_𝑠,𝑡_ is the number of reads for the same genotype in the total population. 𝐶_𝑓_ and 𝐶_𝑡_ denote the number of cells in the fluorescently sorted and total populations, respectively. The enrichment score for variant 𝑠, 𝐸_𝑠_, was then defined as 𝐸_𝑠_ = 𝑝_𝑠,𝑓_ if the sorted population (whichever was smaller in the FACS experiment) represented the bound phenotype, and 𝐸_𝑠_ = 1 − 𝑝_𝑠,𝑓_ otherwise. Standard errors were estimated from replicate experiments.

### Variant-specific fluorescence inference

For SortSeq and TiteSeq experiments, using sequencing and flow cytometry data, we calculated the mean log-fluorescence of each genotype 𝑠 at each concentration 𝑐, as follows:

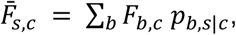

where 𝐹_𝑏,𝑐_ is the mean log-fluorescence of bin 𝑏 at concentration 𝑐, and 𝑝_𝑏,𝑠|𝑐_ is the inferred proportion of cells from genotype *s* that are sorted into bin 𝑏 at concentration 𝑐, which is estimated from the read counts as:

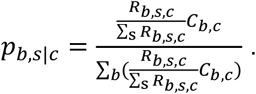

Here, 𝑅_𝑏,𝑠,𝑐_ represents the number of reads from genotype *s* that are found in bin 𝑏 at concentration 𝑐, and 𝐶_𝑏,𝑐_ refers to the number of cells sorted into bin 𝑏 at concentration 𝑐. This log fluorescence is also used as phenotype values for SortSeq measurements.

We then computed the uncertainty for the mean log-fluorescence:

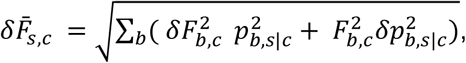

where 𝛿𝐹_𝑏,𝑐_ is the spread of the log fluorescence of cells sorted into bin 𝑏 at concentration 𝑐. The error in 𝑝_𝑏,𝑠|𝑐_ emerges from the sampling error, which can be approximated as a Poisson process, such that:

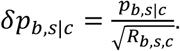

### Binding affinity, K*_D_* inference

For affinity measurements across varying ligand concentrations, we estimated the apparent dissociation constant *K*_D,app_ or each genotype as previously described. Briefly, we fit the logarithmic form of the Hill function to the mean log-fluorescence 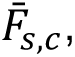 which we had previously computed, as a function of ligand concentration 𝑐:

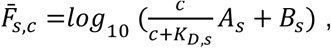

where 𝐴_𝑠_ represents the increase in fluorescence at ligand saturation and 𝐵_𝑠_ denotes the background fluorescence level. The fits were performed using the *curve_fit* function in the Python package *scipy.optimize*. Across all genotypes, we imposed bounds on the values of 𝐴_𝑠_ to be 10²-10⁶, 𝐵_𝑠_ to be 1-10⁵, and *K*_D,s_ to be 10⁻¹⁴-10⁻⁵. For each measurement, the inferred *K*_D,s_ values were averaged across two experimental replicates after excluding fits with poor quality (𝑅^2^ < 0.8 or standard error > 1).

### Background effect comparisons

We performed background comparison analyses to assess how mutation effects varied across different genetic backgrounds (Figure 1C). For Library C, we calculated, for each mutation-background pair (*m*, *g*), the effect of adding mutation *m* across all combinations of other mutations (excluding *m*), with the background *g* treated as part of the combination. For Library M, which was not combinatorially complete, we instead estimated the effect of mutation *m* in background *g* by first computing the average phenotype of genotypes containing *m* and comparing it to the average phenotype of genotypes lacking *m*. The difference between these averages provided the estimated effect of mutation *m* in that background.

### Epistasis analysis

To quantify epistatic interactions in the Library B and Library C datasets, we used the Python package pymochi. Briefly, each inference consists of two concurrent modeling steps. First, an additive model comprises the weights of individual mutations and their interactions. This additive model is then passed to a neural network with two hidden layers to capture nonlinear relationships between the additive biochemical model and the observed fluorescence measurements.

For TiteSeq measurements, the additive biochemical model corresponds to a simple linear combination of mutation effects (mochi transformation: ‘Linear’). Specifically, we assume that the logarithm of the apparent dissociation constant, log 𝐾_D,app_, is proportional to the change in free energy. In contrast, for SortSeq measurements on Library C, we used the ‘ThreeFractionBound’ transformation for each of the two tested concentrations (see Codes). In this case, the same additive weights are jointly inferred across both low and high concentration phenotypes, as well as for the expression phenotype. For both TiteSeq and SortSeq measurements, we reported the inferred additive model weights, including first-and higher order coefficients. The full *K*-order additive biochemical model can be expressed as:

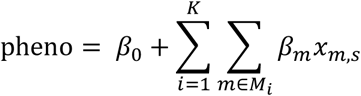

where 𝛽_𝑚_ denotes the coefficient for the combination of mutation 𝑚 (a single-mutation coefficient for 𝑖 = 1 or interaction coefficient otherwise), containing all combinations of *i* mutations.

To determine the optimal value of *K*, we performed 10-fold cross-validation for all *K* ≤ 6. For each *K*, the data were divided into ten subsets, with each serving once as a test set for a model trained on the remaining nine. The optimal *K* was chosen as the value that maximized the average prediction performance (R²) averaged across the ten test sets.

### Statistical analyses and visualization

All data processing and statistical analyses were performed using R^42^ and python^43^. All figures were generated using ggplot2^44^ and matplotlib^45^.

## ACKNOWLEDGEMENTS

TD acknowledges support from the Human Frontiers Science Program Postdoctoral Fellowship. MMD acknowledges support from NSF grant PHY-1914916 and NIH grant GM104239.

