## Supplementary Figures for "Epistasis and background dependence in the evolution of Omicron variants of the SARS-CoV-2 Spike protein"

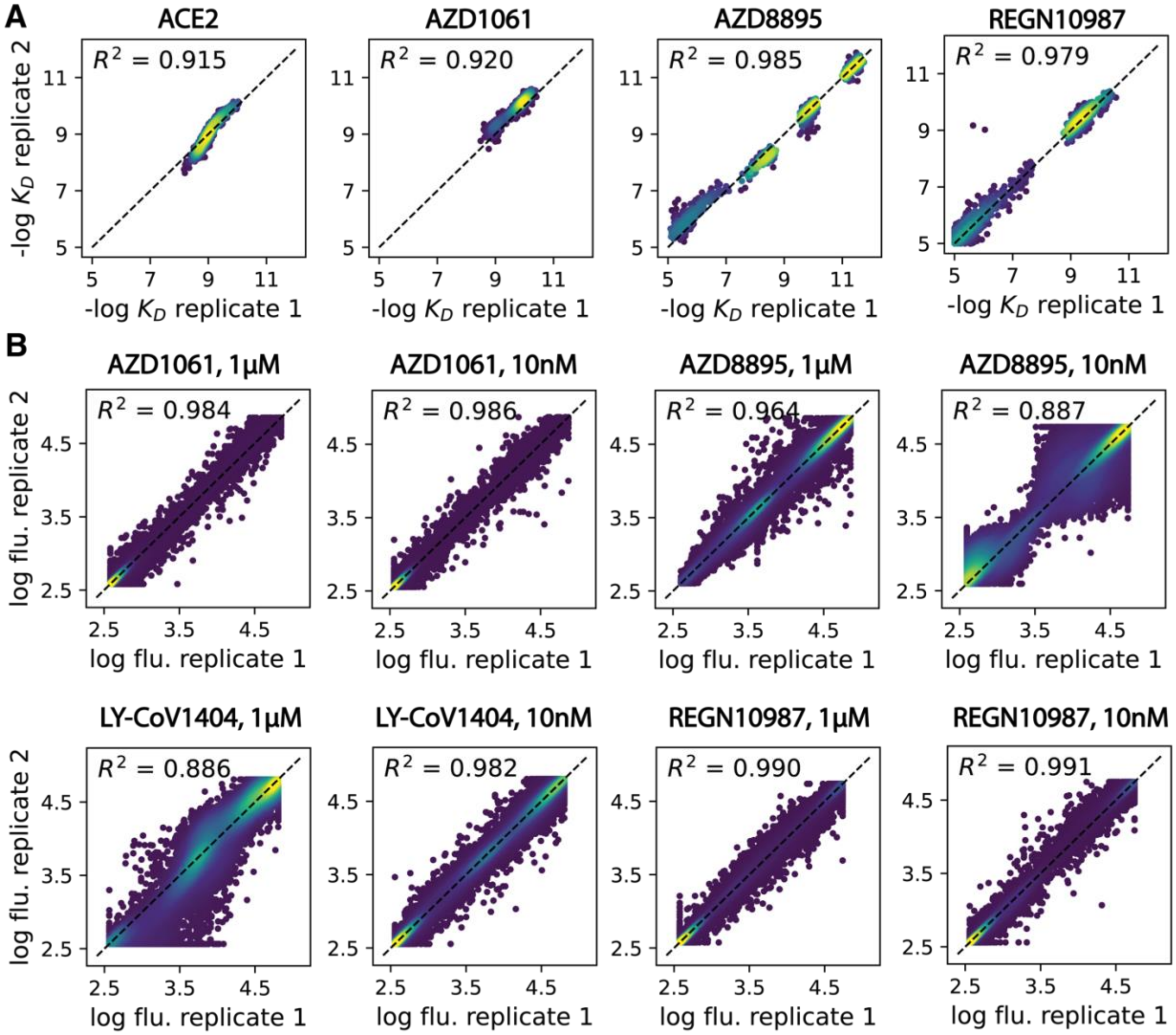

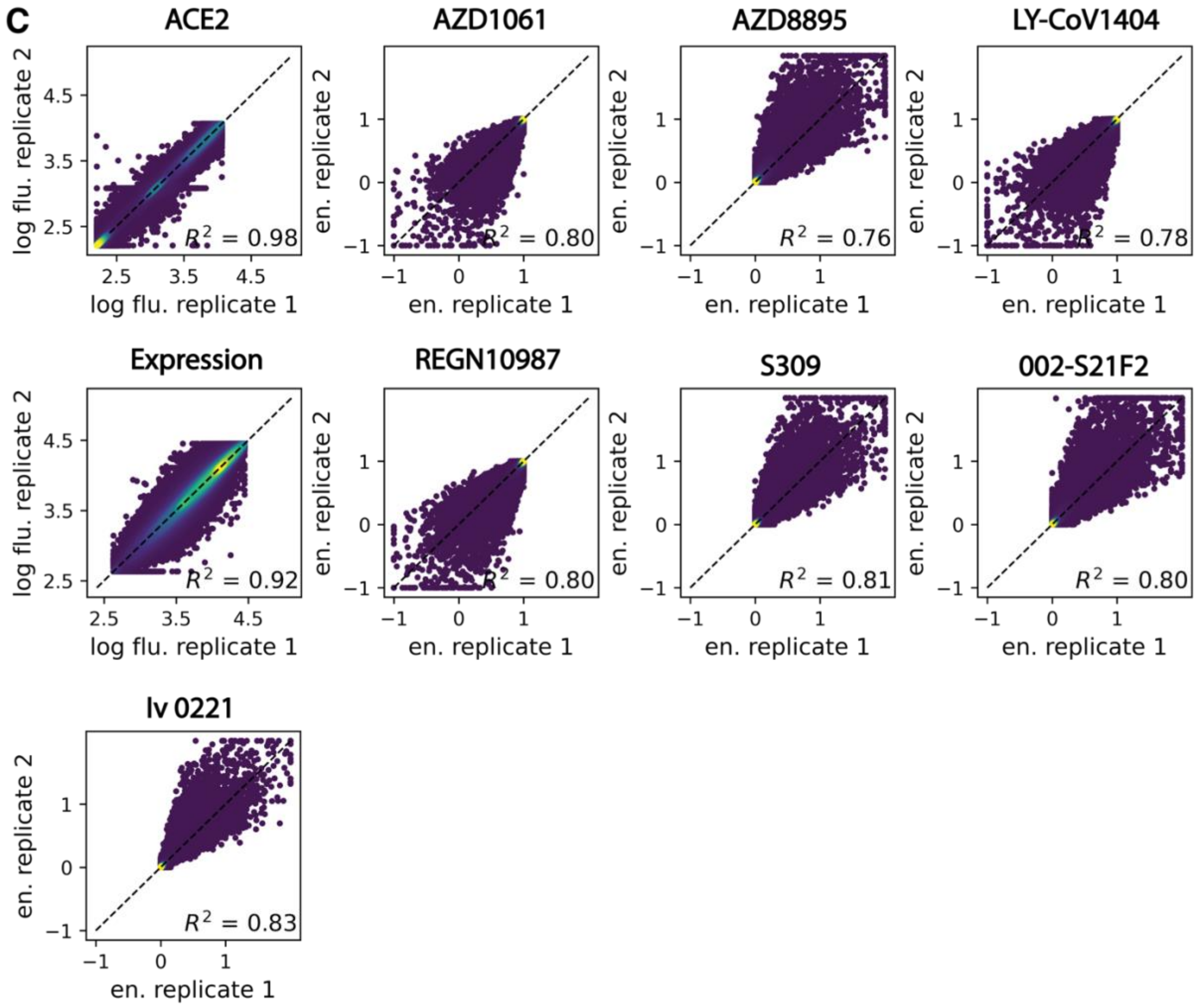


**Supplementary Figure 1: Phenotype correlations between measurement replicates.** Scatter plots showing replicate correlations for each phenotype measured in Library B (**A**), C (**B**), and M (**C**). Spearman correlation coefficient ($R^2$) is shown within each plot.

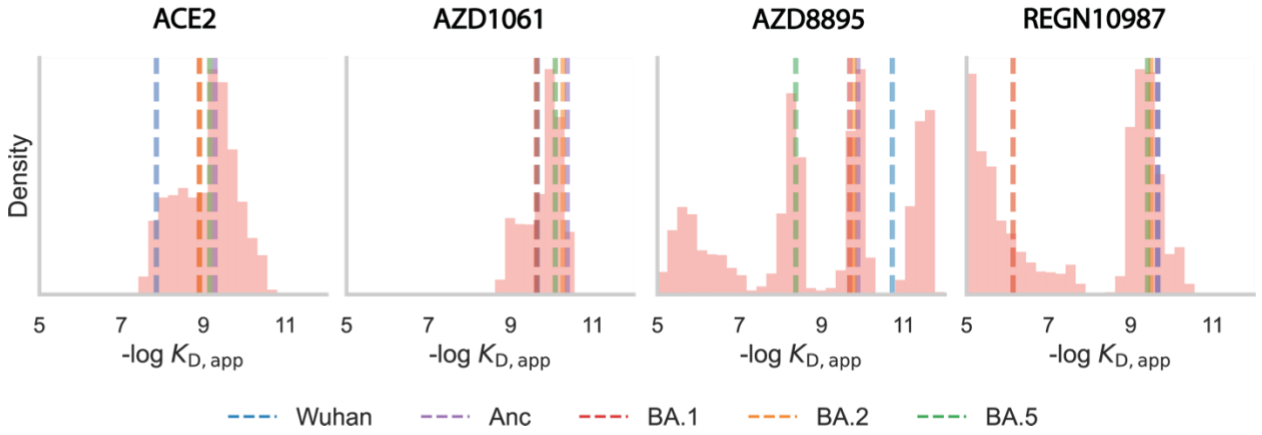


**Supplementary Figure 2: Phenotype distribution for Library B variants.** Histogram of -log $K_{D,app}$ values for variants in the library. Dashed vertical lines, colored according to the variant, indicate values for specific variants of concern.

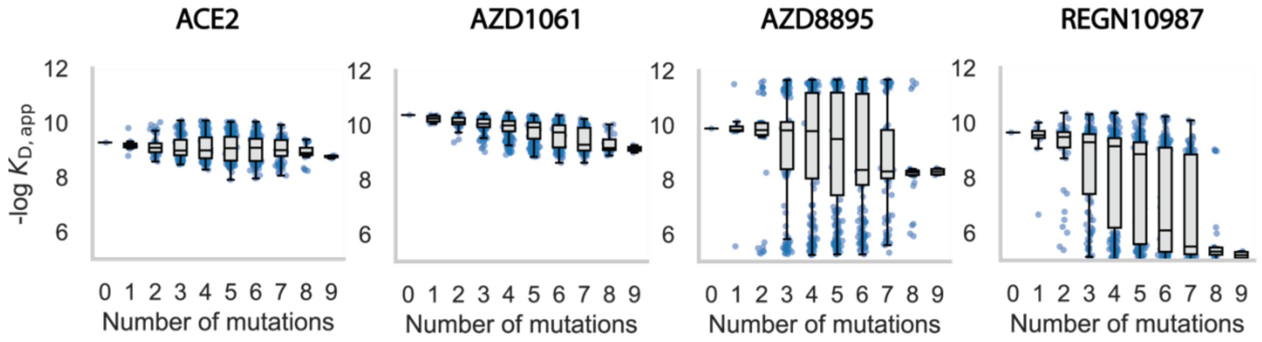


**Supplementary Figure 3: Phenotype distribution for Library B variants grouped by the number of mutations.** Boxplots of -log $K_{D,app}$ values for variants carrying 0 to 9 mutations for each measured phenotype. In each boxplot, the box represents the interquartile range (25th to 75th percentile), the horizontal line inside the box indicates the median, whiskers extend to the most extreme points within 1.5× the interquartile range, and individual outliers are shown as points with jitter.

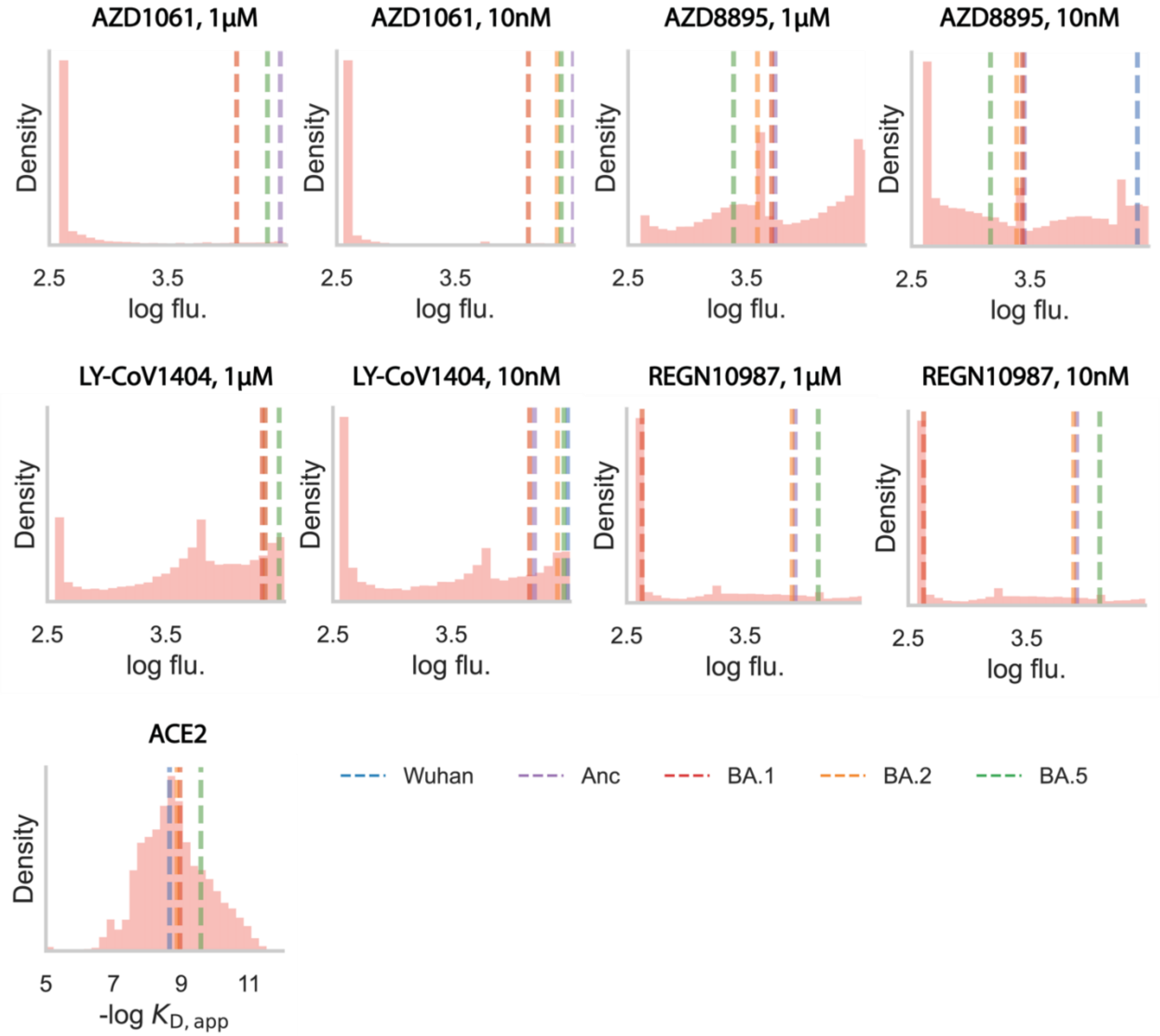


**Supplementary Figure 4: Phenotype distribution for Library C variants.** Histogram of -log $K_{D,app}$ (for ACE2) and log fluorescence (for mAbs) values for variants in the library. Dashed vertical lines, colored according to the variant, indicate values for specific variants of concern.

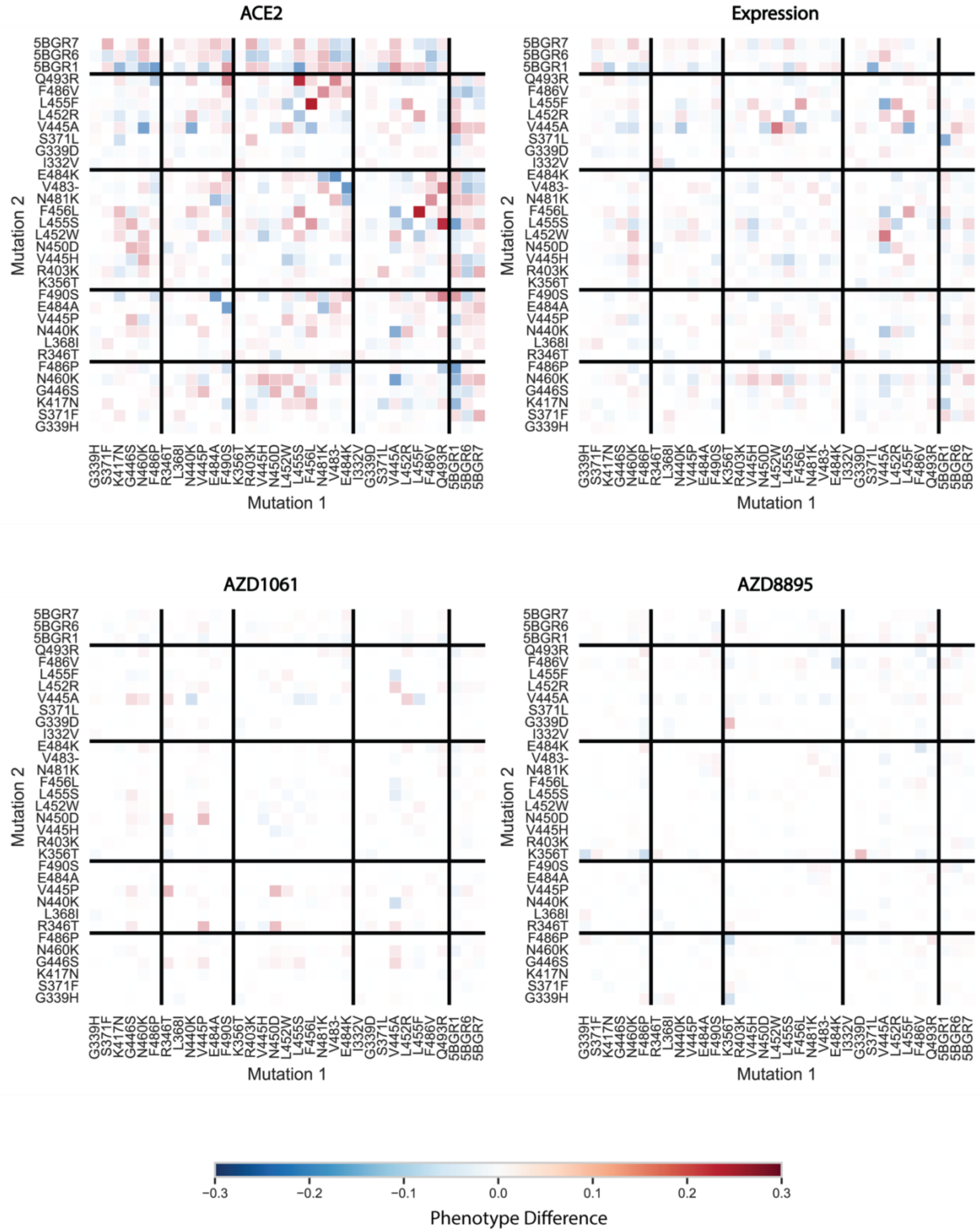
ACE2
Expression
AZD1061
AZD8895
Mutation 2
5BGR7 5BGR6 5BGR1 Q493R F486V L455F L452R V445A S371L G339D I332V E484K V483- N481K F456L L455S L452W N450D V445H R403K K356T F490S E484A V445P N440K L368I R346T F486P N460K G446S K417N S371F G339H
Mutation 1
G339H S371F K417N G446S N460K F486P R346T L368I N440K V445P E484A F490S K356T R403K V445H N450D L452W L455S F456L N481K V483- E484K I332V G339D S371L V445A L452R L455F F486V Q493R 5BGR1 5BGR6 5BGR7
−0.3 −0.2 −0.1 0.0 0.1 0.2 0.3
Phenotype Difference

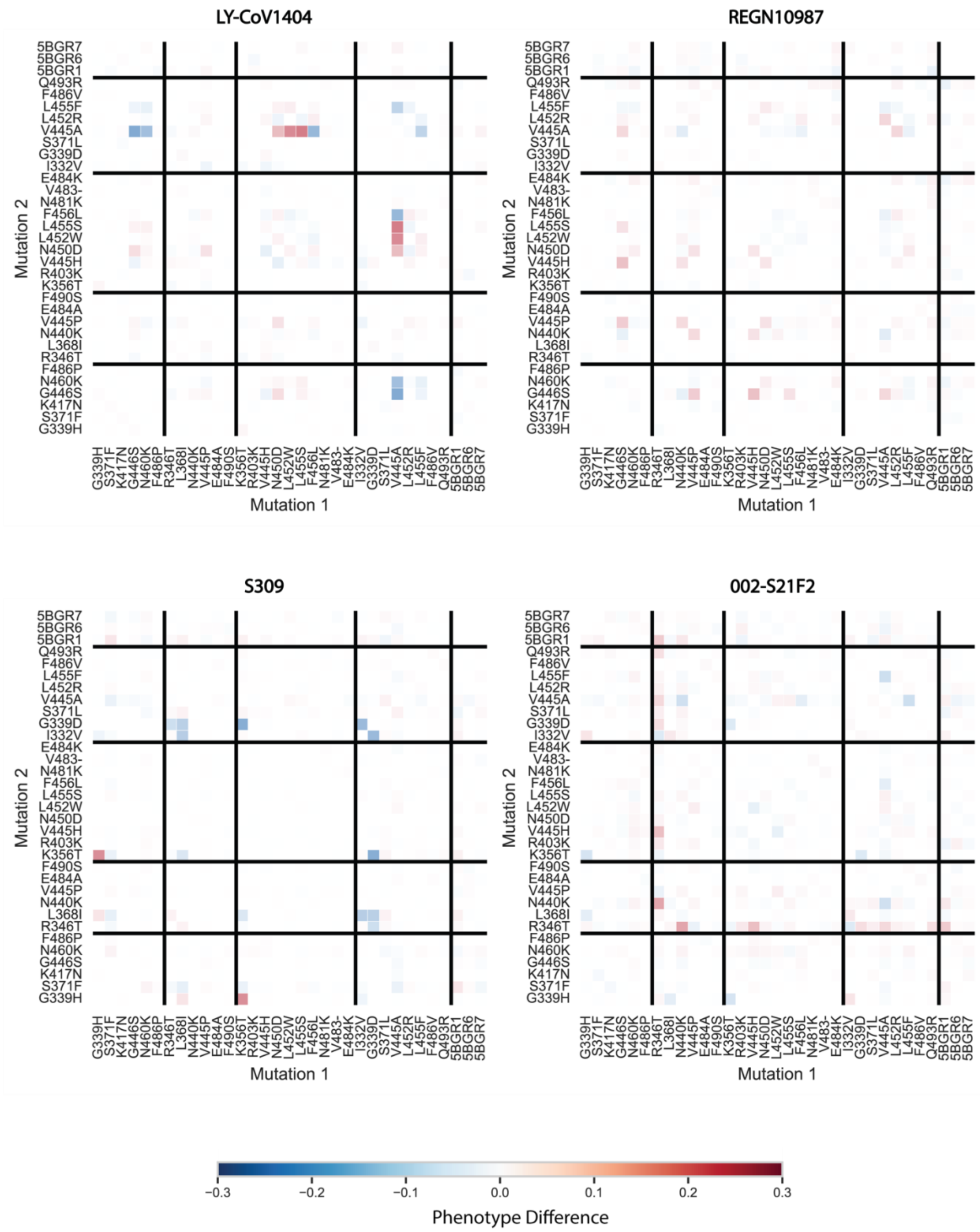

LY-CoV1404
REGN10987
S309
002-S21F2
Mutation 2
5BGR7 5BGR6 5BGR1 Q493R F486V L455F L452R V445A S371L G339D I332V E484K V483- N481K F456L L455S L452W N450D V445H R403K K356T F490S E484A V445P N440K L368I R346T F486P N460K G446S K417N S371F G339H
Mutation 1
G339H S371F K417N G446S N460K F486P R346T L368I N440K V445P E484A F490S K356T R403K V445H N450D L452W L455S F456L N481K V483- E484K I332V G339D S371L V445A L452R L455F F486V Q493R 5BGR1 5BGR6 5BGR7
−0.3
−0.2
−0.1
0.0
0.1
0.2
0.3
Phenotype Difference

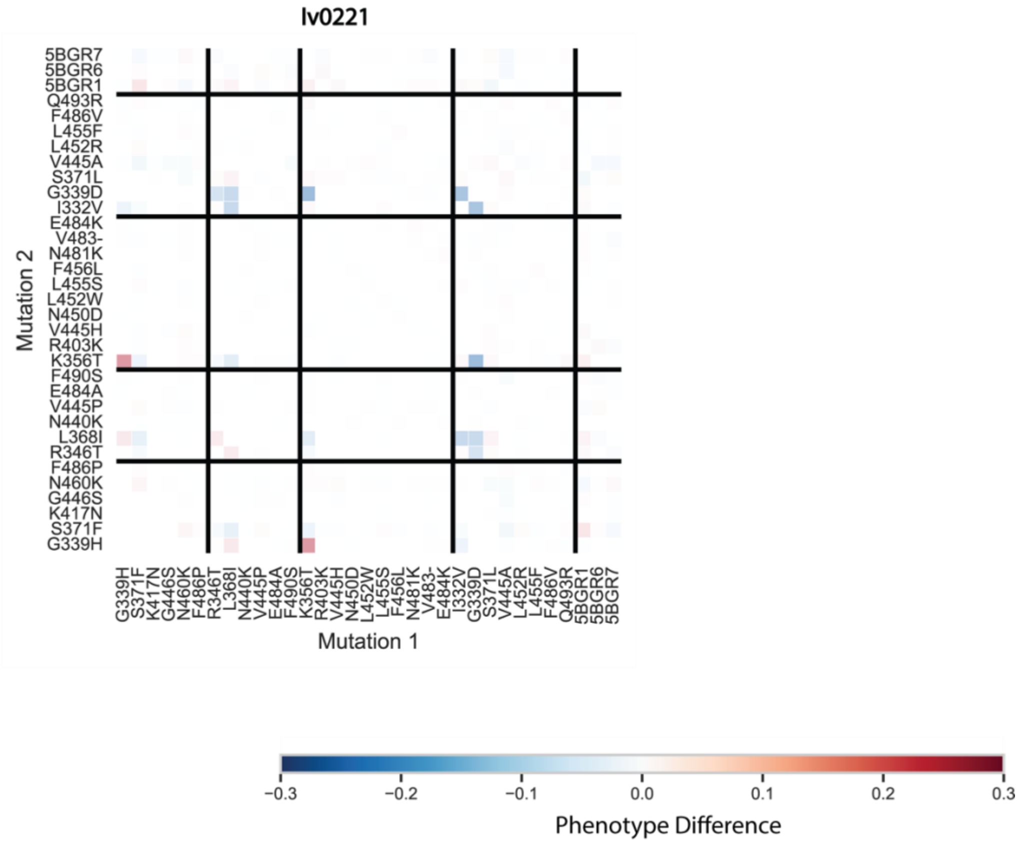


**Supplementary Figure 5: Library M mutation/background interactions across measured phenotypes.** For each phenotype, heatmap (low → high, blue → red) showing pairwise interaction effects between mutations or backgrounds (x and y axes). Thick black lines separate mutation groups, ordered from bottom to top and from left to right, as follows: shared between XBB and JN.1 lineages, specific to XBB.1 lineage, specific to JN.1 lineage, single mutations not in either lineage, and background changes relative to Background 5. Interactions are calculated as the difference between Mutation 2 effects (each computed as in Figure 4) on genotypes carrying Mutation 1 and on genotypes lacking Mutation 1.
